# Using semantic search to find publicly available gene-expression datasets

**DOI:** 10.1101/2025.03.13.643153

**Authors:** Grace S. Brown, James Wengler, Aaron Joyce S. Fabelico, Abigail Muir, Anna Tubbs, Amanda Warren, Alexandra N. Millett, Xinrui Xiang Yu, Paul Pavlidis, Sanja Rogic, Stephen R. Piccolo

**Affiliations:** Department of Biology, Brigham Young University, Provo, Utah, USA; Institute of Biosciences and Technology, Texas A&M Health Science Center, Houston, TX, USA; Department of Psychiatry, University of British Columbia, Vancouver, British Columbia, Canada; Michael Smith Laboratories, University of British Columbia, Vancouver, British Columbia, Canada

## Abstract

Millions of high-throughput, molecular datasets have been shared in public repositories. have been shared in public repositories. Researchers can reuse such data to validate their own findings and explore novel questions. A frequent goal is to find multiple datasets that address similar research topics and to either combine them directly or integrate inferences from them. However, a major challenge is finding relevant datasets due to the vast number of candidates, inconsistencies in their descriptions, and a lack of semantic annotations. This challenge is first among the FAIR principles for scientific data. Here we focus on dataset discovery within Gene Expression Omnibus (GEO), a repository containing 100,000s of data series. GEO supports queries based on keywords, ontology terms, and other annotations. However, reviewing these results is time-consuming and tedious, and it often misses relevant datasets. We hypothesized that language models could address this problem by summarizing dataset descriptions as numeric representations (embeddings). Assuming a researcher has previously found some relevant datasets, we evaluated the potential to find additional relevant datasets. For six human medical conditions, we used 30 models to generate embeddings for datasets that human curators had previously associated with the conditions and identified other datasets with the most similar descriptions. This approach was often, but not always, more effective than GEO’s search engine. Our top-performing models were trained on general corpora, used contrastive-learning strategies, and used relatively large embeddings. Our findings suggest that language models have the potential to improve dataset discovery, perhaps in combination with existing search tools.

## Introduction

Publishers and funding agencies often mandate that researchers deposit data in public repositories so that others can verify their findings and build on their work. However, making data *accessible* on the Internet is not enough. To accelerate progress for the research community as a whole—and to ensure that funding agencies maximize their investments—data must also be *findable*, *interoperable*, and *reusable*^1^. Here we focus on the first of these requirements: the ability for researchers to find datasets that have been shared publicly. Without this capability, the benefits of interoperability and reusability are reduced.

As an example of a large, highly used, public repository, we focus on Gene Expression Omnibus (GEO)^2^, which houses data from more than 7,500,000 biological samples, spanning more than 240,000 data series. (In this paper, we use the term *series* interchangeably with *dataset* because the latter reflects the parlance commonly used to describe data-finding efforts, even though GEO also contains “DataSets,” which are curated series.) GEO contains many types of high-throughput molecular data; transcriptomic data are the most common. Similar to other repositories, GEO provides search capabilities. Users enter keywords, which are matched against metadata associated with samples or series. These metadata include the title, description, profiling platform, and descriptions of sample attributes. Additionally, for many studies, metadata include Medical Subject Heading (MeSH) terms, which are human curated^3^. After performing a search, users can filter results based on the species name, assay type, sample size, and other criteria. Despite the flexibility of this search tool, in our experience, these searches often yield irrelevant datasets and miss relevant ones, thus requiring time- and labor-intensive reviews and additional searches.

As one way to address this problem, researchers unaffiliated with GEO have curated lists of transcriptomic datasets on particular research topics^4–9^. Others have annotated GEO series with labels describing their biomedical context^10–14^. One example of the latter is Gemma^10^. Its curators have manually reviewed transcriptomic data series (mostly from GEO) and annotated them. Gemma focuses primarily on human, mouse, and rat datasets, especially those related to the nervous system. Gemma users can filter series based on these labels. However, due to the manual effort required to assign such labels—and Gemma’s focus on particular types of datasets—this resource houses fewer than 10% of all GEO series. Furthermore, the annotation process is subjective. Labels might be inconsistent from one study to the next^15^; other domain experts might disagree with the labels; the labels might not be comprehensive; and the labeling efforts might be limited to particular research areas.

To facilitate dataset finding for GEO and other repositories, third-party researchers have created alternative ways to search for transcriptomic datasets (and other types of data). For example, *Omics Discovery Index* and *DataMed* provide free-text search capabilities, along with filters for publication year, species, data type, source repository, tissue type, and technology type^16–18^.

These tools support searching for datasets in GEO and other repositories. However, like GEO, their search methodologies primarily rely on keyword expansion and matching. As a result, they may struggle to account for semantic context effectively and may not handle polysemy (words with multiple meanings), synonyms, and different word forms. As an alternative, some researchers have used manual curation, natural language processing (NLP), and/or data inferences to aid in identifying series or samples based on species, cell type, tissue type, disease type, and/or assay type^19–25^. Hawkins, et al. used neural-network models to create an embedding—a numeric representation of words and phrases—for each sample^20^. Such approaches are capable of representing data semantics, even when words are misspelled or when different terms are used to describe the same concepts^26^. However, their approach focused on identifying and characterizing samples, whereas our goal is to support finding at the series level where longer-form descriptions are available. PEPHub uses a language model to support finding GEO series, but they did not report benchmarks against other techniques^27^. Patra, et al. developed a system for recommending GEO series based on embeddings of research articles that a scientist had published previously^28^. However, a researcher’s past publications may not coincide directly with future interests, and this approach may not be effective for early-career researchers.

For this study, we assume that a researcher has used existing tool(s) to find GEO series on a given research topic and wishes to find additional ones. We hypothesized that we would be able to detect semantically similar datasets based on metadata. We focus on methodologies that summarize text as embeddings. In recent years, computational researchers have developed diverse methodologies for training embeddings. Early examples include Word2Vec and GloVe^29–31^. These algorithms train neural networks on large text corpora. Given a particular word (or subword), they characterize the context in which that (sub)word is used, based on surrounding (sub)words, in a context window. More recently, BERT and transformer-based models have become widely used for NLP^31,32^. The attention mechanisms of these techniques account for relationships between a given (sub)word and all words in a given sentence, regardless of their distance from the (sub)word. Researchers have trained such models using general corpora like Wikipedia, books, news articles, and other sources. Others have trained models using domain-specific corpora, including some specific to biology and/or medicine^33–37^. Furthermore, it is possible to build on pre-trained models using a process known as fine tuning.

With this approach, a researcher uses an existing model—which might have been trained on billions of examples—and performs additional training with domain-specific examples. This strategy yields significant cost and energy savings compared to training a model from scratch and may yield better results for NLP tasks in that domain.

Prior studies have shown that relatively large corpora are often better than smaller ones^38^ and that training on domain-specific corpora is often better than using general corpora^35,39^. However, the fast pace with which this field is changing necessitates additional research. We performed a benchmark study, starting with a simple, word-counting approach and then evaluating 30 language models ranging in complexity from continuous bag-of-words models to transformer-based models. We applied these models in the context of GEO dataset finding, in which researchers have provided ad hoc descriptions that differ in their length, semantics, and level of detail. One challenge is that, in a typical scenario, the number of datasets relevant to a particular research topic is small (perhaps between 5 and 100) compared to the total number of GEO datasets (100,000s). Therefore, we have evaluated the effects of this imbalance. Our focus is not necessarily on identifying particular model(s) or optimization techniques that outperform all others. Instead, our goal is to shed light on the extent to which modern language models have the potential to aid with dataset discovery.

## Methods

### Annotated data collection

On April 18, 2024, we used geoFetch (version 0.12.5)^40^ and custom Python code (https://python.org, version 3.12) to retrieve metadata associated with all available GEO series. We retained series that matched the following criteria:

- Were *not* marked as “retired.”
- Were listed as coming from *homo sapiens* samples.
- Were associated with the “expression profiling by array” experiment type and used an Affymetrix, Illumina, or Agilent platform OR were associated with the “expression profiling by high throughput sequencing” experiment type and used an Illumina platform.
- Were *not* a GEO SubSeries. We excluded these to avoid bias in our machine-learning analysis (having identical descriptions in the reference and comparison groups) and because it would be easy for researchers to identify a SubSeries if they have identified its associated SuperSeries.

A total of 48,893 series matched these criteria. For each of these series, we use diverse models to generate numeric vectors (embeddings) that summarized the text describing the series. When creating these vectors, we used the title, summary, and overall design (when available) as inputs. Before doing so, we converted the text to lower case and removed URLs, non-alphanumeric characters, and extra white space; we also removed HTML tags using the Beautiful Soup (version 0.0.2) package^41^. Because the *fastText* and *word overlap* models (described below) do not rely on (and may be distracted by) stop words when inferring semantic meaning, we removed stop words, as defined by the *nltk* package (version 3.8.1)^42^. For the other models—most of which are transformer-based—we retained stop words because those models capture the semantic context of words based on the entire sentence or document. When text inputs were longer than 256 characters, we sometimes (details below) split the text into chunks, allowing for 20 characters of overlap between chunks. We chose 256 characters as the threshold because the smallest embedding size was 300 characters and because some models truncate inputs of approximately this size.

On April 19, 2024, we queried Gemma^10^ to identify all GEO series that had been annotated in that resource and that overlapped with the 48,893 we had identified (n = 5,997). We also selected series that human curators had associated with one of six medical conditions: juvenile idiopathic arthritis, triple negative breast carcinoma, Down syndrome, bipolar disorder, Parkinson’s disease, and neuroblastoma. Some of these series coincide with our research interest in cancer; others coincide with Gemma’s focus on neurological conditions. Together, these series reflect a range of sample sizes (some conditions have been more highly studied than others).

### GEO Advanced Search Builder queries

On May 8, 2024, we used GEO’s Advanced Search Builder to identify datasets (series) associated with each of the six medical conditions. We attempted three types of queries using:

1) The name of the medical condition as a keyphrase.
2) The name of the medical condition plus synonyms for the medical condition as keyphrases. The synonyms were defined by the Mondo Disease Ontology^43^, which we queried using BioPortal^44^.
3) The MeSH Subject Heading term^45^ for the condition.

For #1 and #2, GEO sometimes expanded our queries by mapping them automatically to MeSH terms. For #3, the MeSH Browser showed the term *Arthritis, Juvenile* (D001171), but this option (or other variations on it) did not appear in GEO. Therefore, we used the broader MeSH term *Arthritis* (D001168). Similarly, GEO did not provide an option for triple-negative breast cancer, even though the MeSH Browser showed a term for *Triple Negative Breast Neoplasms* (D064726). Therefore, we searched using the broader term *Breast Neoplasms* (D001943). Our source-code repository contains files with the exact queries that we used and the results of each query.

### Experimental design

For each medical condition, we identified all datasets annotated for that condition in Gemma. We randomly assigned half of these to a “reference set” and half to a “comparison set A” (Figure 1). When there was an even number of datasets, the reference set and comparison set A were the same size. When there was an odd number of datasets, the reference set contained one more dataset than comparison set A. From the remaining Gemma datasets (including those from other medical conditions), we created “comparison set B”. Using each medical condition’s reference set, we averaged the embeddings from the individual datasets and used the cosine-similarity metric to calculate the distance between this embedding and each dataset in comparison sets A and B. We used cosine similarity because it is easily calculated, interpretable, invariant to the magnitude of vectors being compared, and widely used in NLP studies. Finally, we ranked the datasets in the comparison sets based on cosine similarity, with the expectation that more effective language models would rank datasets from comparison set A higher than those from comparison set B.

**Figure 1:**
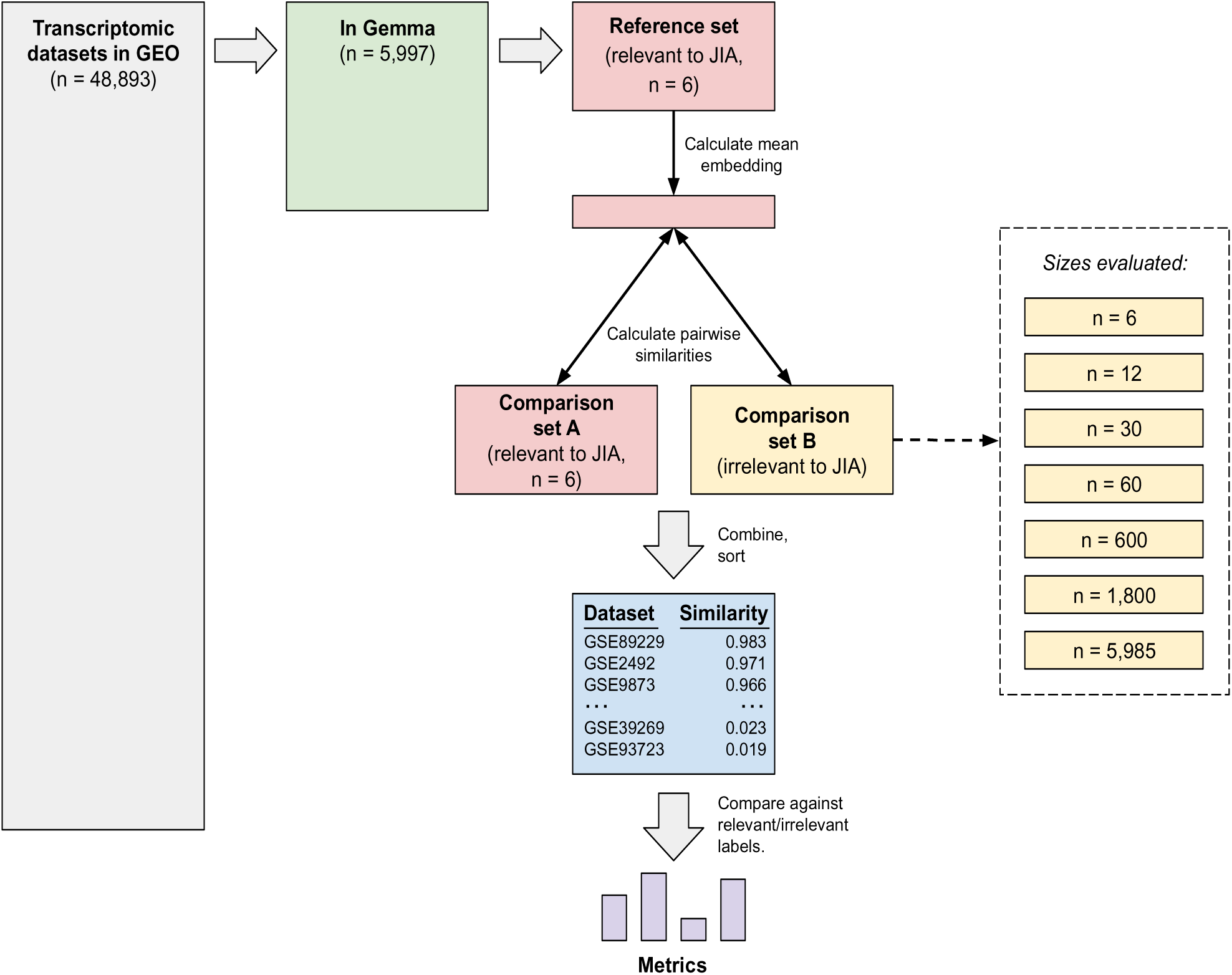
Evaluation process for comparing language models’ abilities to identify relevant datasets. This diagram illustrates the process of evaluating the language models’ performance for a particular medical condition. Here we use juvenile idiopathic arthritis (JIA) as an example. After identifying transcriptomic datasets in GEO, we selected those that had been annotated in Gemma as relevant to the medical condition. We randomly divided these datasets into two evenly sized groups: a reference set and comparison set A. We assigned datasets not annotated as relevant to the medical condition to comparison set B. We varied the size of comparison set B to provide insight about the effects of imbalance between the numbers of relevant and irrelevant datasets. Using a given language model, we generated embeddings for datasets in the reference set and both comparison sets. We calculated the mean embedding for the reference set and then calculated the cosine similarity between this mean embedding and the embedding for each dataset in comparison sets A and B. Finally, we calculated metrics to quantify the extent to which each language model ranked relevant datasets higher than irrelevant ones. (Boxes are not necessarily drawn to scale.)

For the language model that performed best according to the Gemma annotations, we performed a second round of validation. For each medical condition, we calculated an average embedding across *all* datasets annotated for that condition in Gemma. Next, we calculated the pairwise cosine similarity between the average embedding and each human transcriptomic dataset *not* in Gemma. Two curators (authors ANM and XXY) reviewed the top-ranked datasets to assess biomedical relevance. As a preliminary step, we selected five datasets for each medical condition from the datasets ranked 51st through 55th and used these for a pilot analysis. Additionally, we included 5 datasets as distractors (not relevant to any of the six medical conditions). Prior to their reviewing these datasets, we created a rubric for the curators (Additional Data File 1). We then asked each curator to independently review the title, summary, and overall-design descriptions and record whether each dataset was relevant to any of the conditions. The curators’ conclusions were identical for all datasets in the pilot.

We randomly divided the top-50 ranked datasets for each medical condition, assigning half uniquely to each curator while assigning 5 to both curators to facilitate an inter-rater agreement evaluation. (Later, we found that three datasets had been selected twice for a given curator, so we removed the duplicate copies.) We created a spreadsheet with information about these datasets for each curator, removed the medical-condition labels, and asked the curators to indicate which medical condition they believed was most relevant to each dataset. Although we did not include distractors in this validation step, we provided an option for the curators to indicate that a given dataset was not relevant to *any* of the six medical conditions. As with the pilot, the curators were blind to the rankings made by the language model. Finally, we repeated this process for the same number of top-ranked datasets identified via GEO’s Advanced Search Builder. For the language model, these conclusions agreed in 54 cases (87.1%). For GEO’s Advanced Search Builder, the conclusions agreed in 55 cases (88.7%).

### Language models

Table S1 provides a summary of the language models we evaluated. For most models, we used the implementation in the Hugging Face repository and the *transformers* package (version 4.37.2)^46^. For other models, we used the *fasttext* (version 0.9.2)^47,48^ or *openai* (1.12.0)^49^ packages. In Table S1, we categorize each model based on the type of corpus on which it was trained (general purpose, biomedical, or scientific defined more broadly) and the high-level methodology used for training, as described in Hugging Face. It also describes any fine tuning that was performed. In addition to these models, we implemented a word-counting method; after identifying all unique words, we quantified similarity as the number of overlapping words between dataset descriptions.

## Data and code availability

All data and code used to perform this analysis have been deposited in a GitHub repository, which is publicly available and uses the MIT License (https://github.com/srp33/GEO_NLP). We used the Python programming language (version 3, https://python.org) and the R statistical software (version 4.4.1) for the analysis. Additionally, we used the *tidyverse* to process and visualize data^50^. To facilitate reproducibility, we executed the analyses within a Docker container^51^.

## Results

We assume that a researcher has found *some* transcriptomic datasets relevant to a given research topic and wishes to find more datasets relevant to the same topic. They might have found datasets previously using a literature search, GEO’s search tools, or some other means. To facilitate finding additional datasets, we used various techniques to independently summarize the descriptions of each dataset as a numeric vector or embedding. These techniques included a simple word-counting method and 30 distinct language models. To compare these approaches, we identified six medical conditions for which annotated datasets were available in both GEO and Gemma: juvenile idiopathic arthritis (n = 12), triple negative breast carcinoma (n = 24), Down syndrome (n = 30), bipolar disorder (n = 34), Parkinson’s disease (n = 109), and neuroblastoma (n = 121). For each medical condition, we randomly divided the datasets into a *reference set* and a *comparison set A*; the remaining datasets that were available in both GEO and Gemma served as “comparison set B” (Figure 1). After averaging the embeddings in the reference set, we calculated the cosine similarity between the average embedding and each dataset in the comparison sets. For each model, we used the area under the precision-recall curve (AUPRC) to quantify the extent to which the model ranked the datasets from *comparison set A* higher than the datasets from *comparison set B*.

The AUPRC varied significantly across medical conditions (Kruskal-Wallis test, p < 0.001; Figure 2). Median performance was highest for Parkinson’s disease and Down syndrome, while it was lowest for triple-negative breast carcinoma and neuroblastoma. The AUPRC also varied significantly across the models (p < 0.001; Figure 3). Ten of the models consistently performed poorly, rarely attaining AUPRC values higher than 0.10. These included the continuous bag-of-words (CBOW) models, which are relatively simple, as well as some transformer-based models that were trained using biomedicine-specific corpora and/or that used large embedding sizes. In all cases except one, these poor-performing models scored lower than the simple word-counting technique. After excluding the word-counting method, we found that the median AUPRC was significantly correlated with embedding size (Spearman’s test, rho = 0.51, p = 0.004) but did *not* differ significantly by corpora type (Mann-Whitney U test, p = 0.46; Figure 4). We also assessed the effects of text chunking. When input texts are relatively large, practitioners sometimes split text into smaller chunks, create an embedding for each chunk, and then average the embeddings. We tried this technique and compared it against the performance we attained without chunking. We found that *not* chunking nearly always performed *better* than chunking (Table S2).

**Figure 2:**
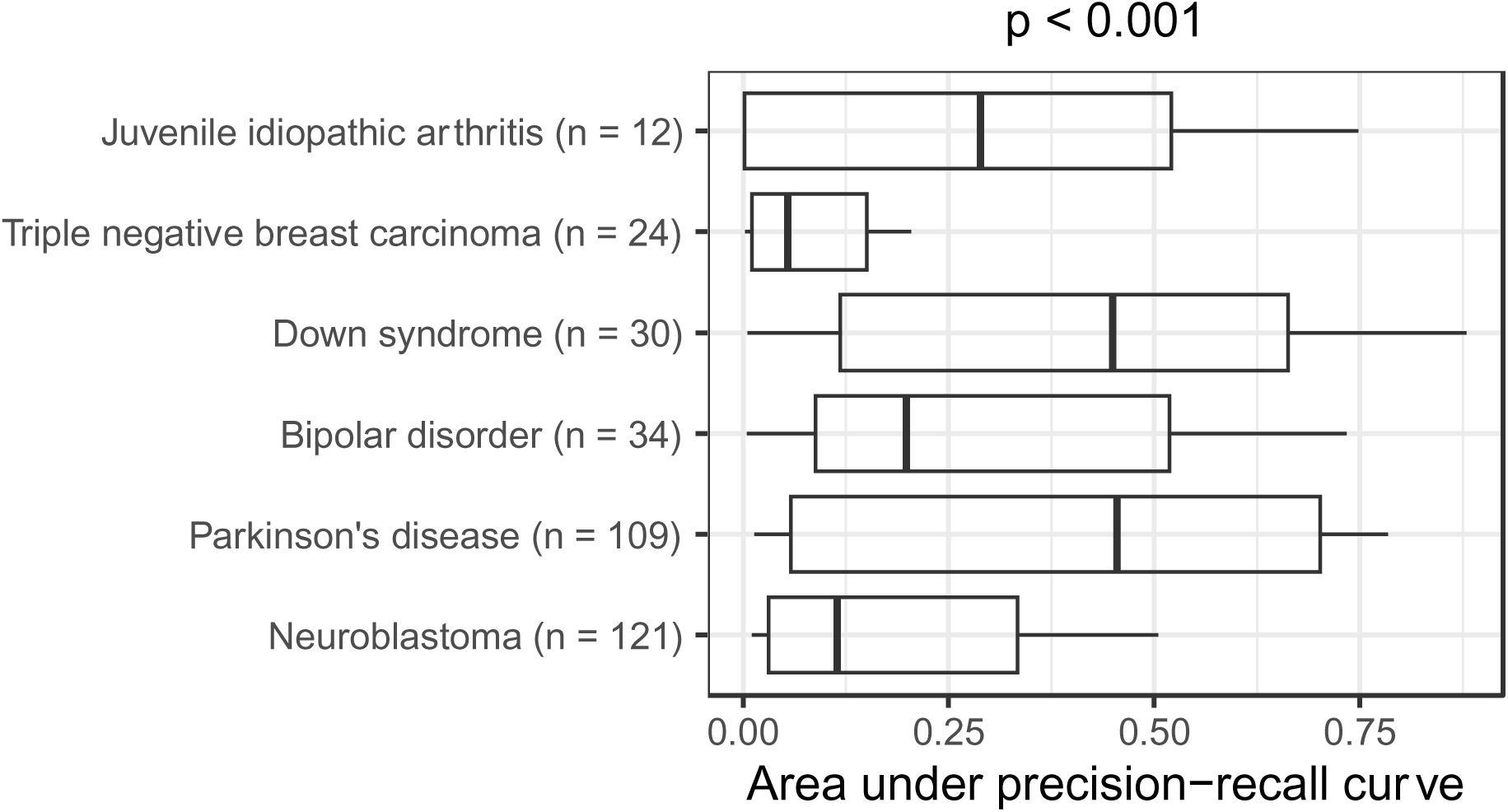
Within-Gemma performance of language models by medical condition. We used language models to identify Gemma datasets that had been annotated as relevant to six medical conditions. The area under the precision-recall curve differed significantly across the conditions. The p-value was calculated using the Kruskal-Wallis test.

**Figure 3:**
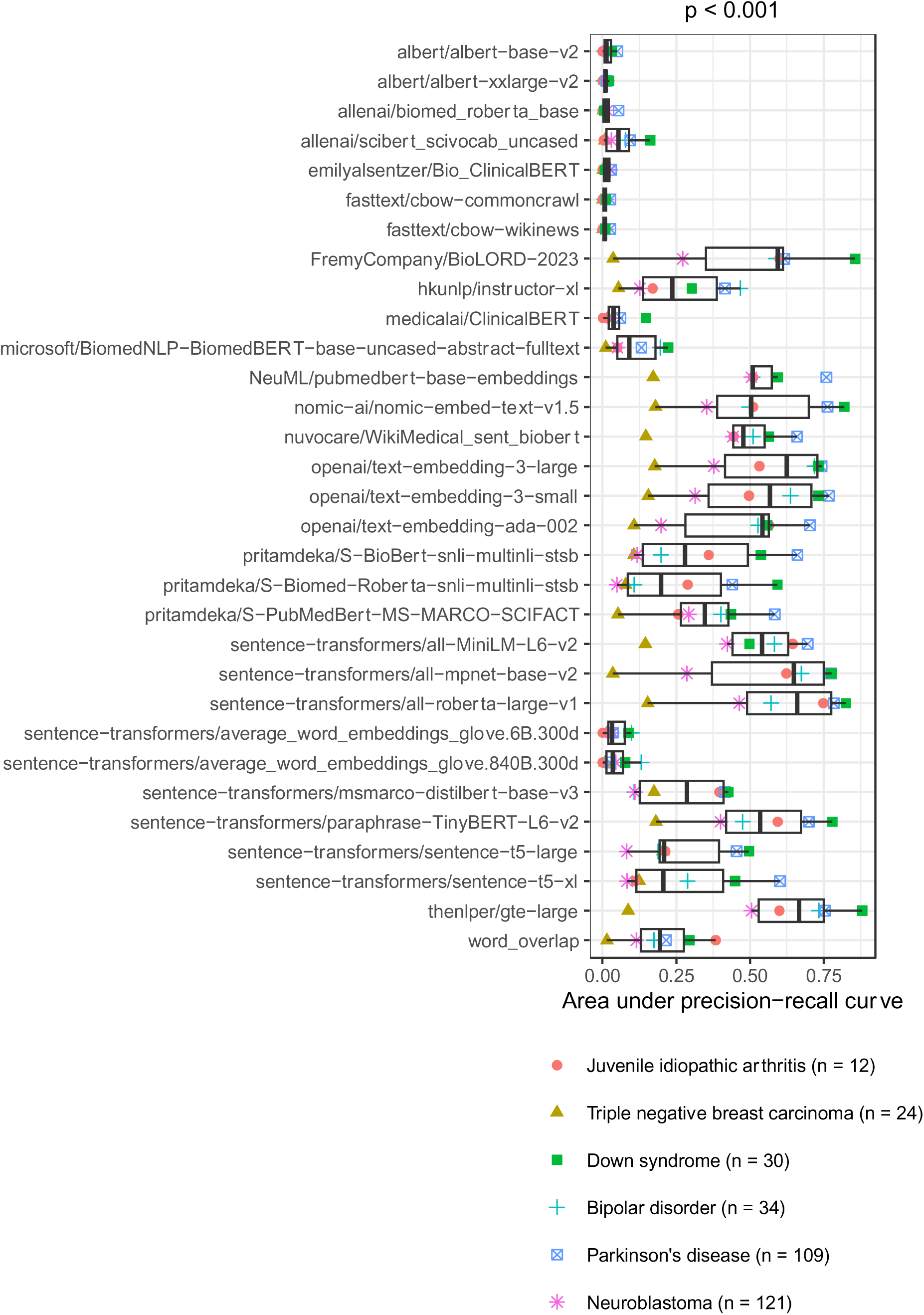
Within-Gemma performance of language models by model. We used language models to identify Gemma datasets that had been annotated as relevant to six medical conditions. The area under the precision-recall curve differed significantly across the models. The p-value was calculated using the Kruskal-Wallis test.

**Figure 4:**
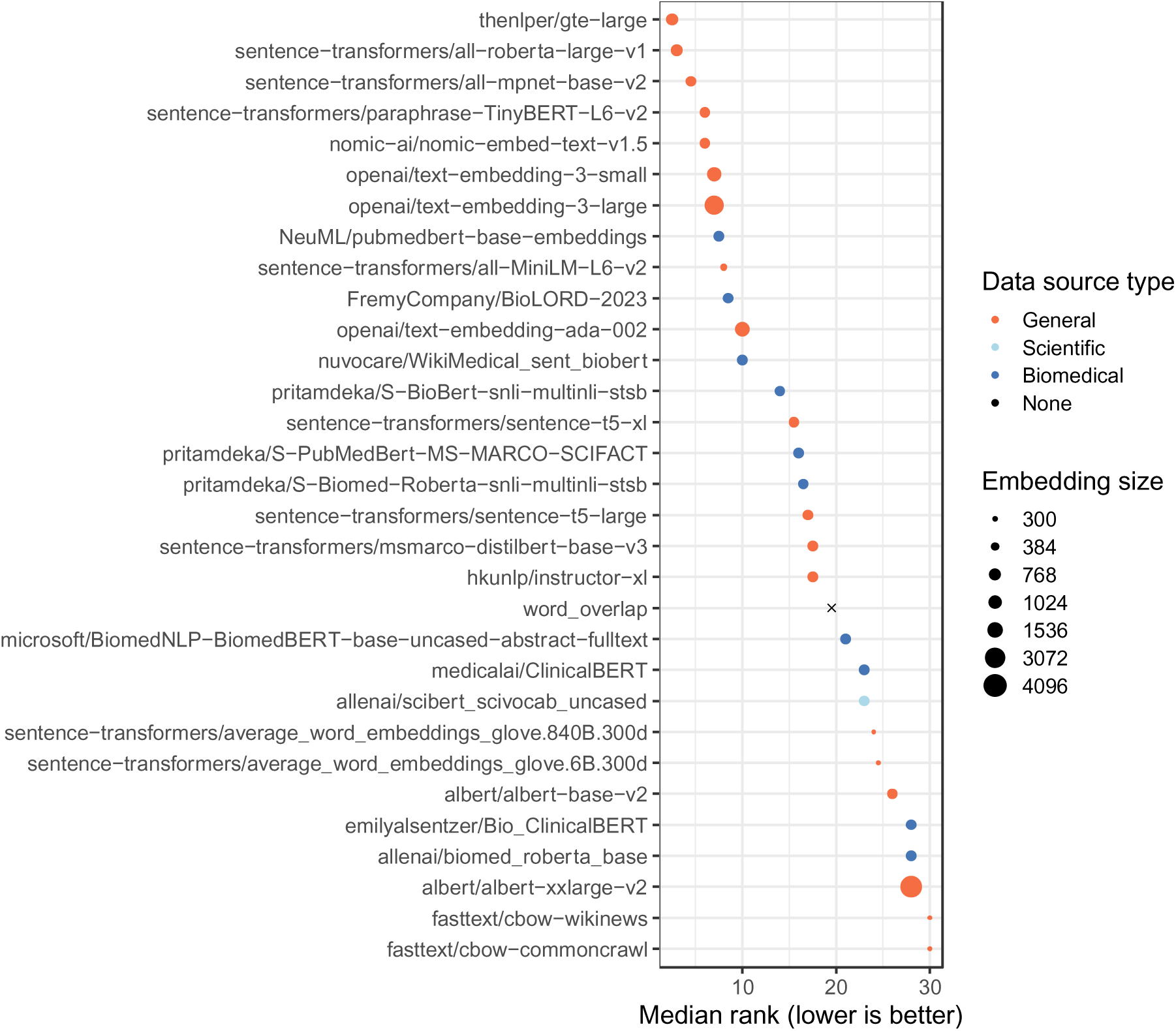
Within-Gemma performance of language models by model, ranked. We used language models to identify Gemma datasets that had been annotated as relevant to six medical conditions. For each medical condition, we ranked the models and calculated the median rank across the conditions. Models with relatively low ranks performed better overall than models with relatively high ranks.

Across the six medical conditions, the top-performing models were `thenlper/gte-large`, `sentence-transformers/all-roberta-large-v1` and `sentence-transformers/all-mpnet-base-v2`. All three were trained using general corpora (not science- or biomedicine-specific) and used moderately sized embeddings (768 - 1024 dimensions). According to the Hugging Face documentation, `thenlper/gte-large` uses multi-stage contrastive learning and was trained by the Alibaba DAMO Academy. It uses the Bidirectional Encoder Representations from Transformers (BERT) framework^31^ and has performed well in the Massive Text Embedding Benchmark^52^. The `sentence-transformers/all-roberta-large-v1` model builds on a pre-trained Roberta model^53^ and was fine-tuned on a dataset with one billion sentence pairs. It uses a self-supervised contrastive learning strategy. The `sentence-transformers/all-mpnet-base-v2` model also builds on a pretrained model, was fine-tuned on one billion sentence pairs, and uses self-supervised contrastive learning. All three models are popular on Hugging Face. For example, in July 2024, each had been downloaded at least 672,000 times. Among other intended uses, all three models are described as being useful for sentence-similarity tasks. We compared the language models’ performance against what was returned using GEO’s Advanced Search Builder (in three variations, see Methods). We used recall (sensitivity) as a metric to quantify the number of search results that needed to be returned before *n* relevant datasets were identified. For juvenile idiopathic arthritis, Down syndrome, and bipolar disorder, at least one of the top-performing models identified all relevant datasets within the top-100 results (Figure S1). All of the top-performing models attained a recall of at least 0.75 for the top-1000 datasets.

Furthermore, the language models nearly always outperformed Advanced Search Builder (Figure 5). One exception was triple-negative breast carcinoma, in which Advanced Search Builder sometimes performed better than the top-performing language models, particularly when using the high-level MeSH term *Breast Neoplasms*.

**Figure 5:**
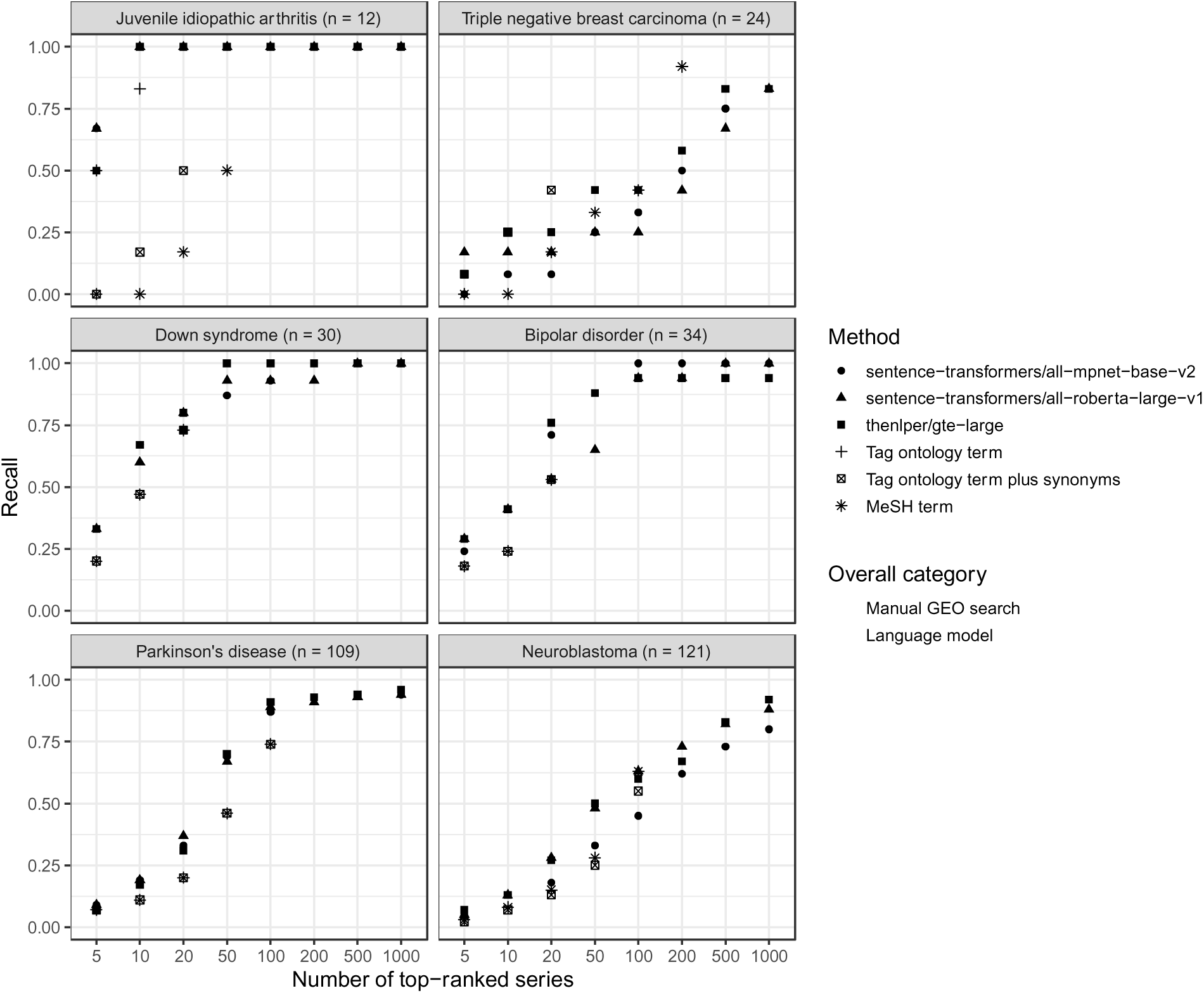
Comparison between manual GEO searches and language-model rankings for within-Gemma series. After manually querying GEO and using language models to identify Gemma series that had been annotated as relevant to six medical conditions, we compared the top-*n* results returned using either approach. In many cases, the GEO search tool returned fewer than 1,000 datasets; therefore, this graph depicts the maximum *n* value that fell below the total number of GEO results for a given medical condition. A recall value of 1.0 indicates that all Gemma-annotated datasets were included in the top-*n* results.

One challenge of dataset finding in this context is that relevant datasets are rare. For example, in our evaluations based on Gemma annotations, we had metadata for 5,997 GEO series, but only 12 of these had been annotated as relevant to juvenile idiopathic arthritis. After splitting the data into reference and comparison sets, we were searching for 6 series among 5,985 candidates (an approximately 1000-to-1 ratio). This imbalance would only increase when searching all GEO series, including those not in Gemma. Therefore, we evaluated the extent to which different levels of imbalance would affect our findings. For juvenile idiopathic arthritis, the AUPRC remained constant for the top-performing models when the ratio was 300-to-1 or smaller.

However, the performance dropped considerably when all Gemma candidates were included (Figure 6). We observed similar patterns for other medical conditions, with the greatest reduction in performance for triple-negative breast carcinoma.

**Figure 6:**
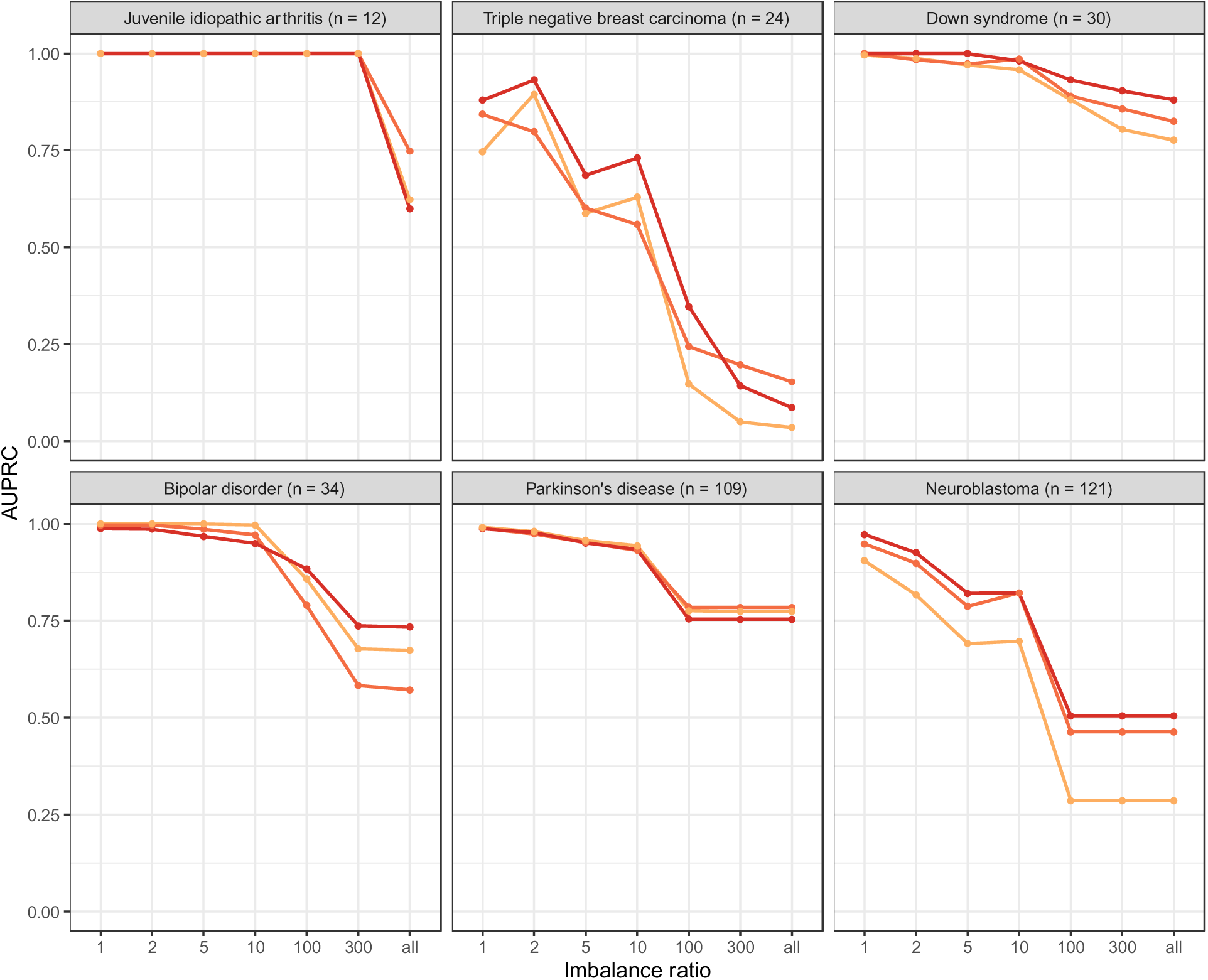
Within-Gemma performance of language models for different levels of imbalance between irrelevant and relevant data series. We used language models to identify Gemma datasets that had been annotated as relevant to six medical conditions. For each medical condition, we performed simulations with increasing levels of imbalance between irrelevant and relevant data series. An imbalance ratio of 1 means that the “other” set had the same number of series as the test set. An imbalance ratio of 10 means that the “other” set had 10 times as many series as the test set. When the imbalance ratio was “all,” we included all available series in the “other” set that were not part of the training or test sets. The lines with color represent the top-3 performing models overall.

To gain additional insight into the top-performing model’s performance (`thenlper/gte-large`), we compared the length (number of characters after cleaning steps) of the dataset descriptions versus the cosine similarity scores for the GEO series in our comparison sets. We surmised that datasets with longer descriptions might be more often ranked as relevant. However, we did *not* observe a significant correlation between these variables (Figure S2).

One ancillary application of our approach is to identify Gemma series that lack relevant annotations. We identified the 25 GEO series that were most ranked highly (across all models) as relevant to a given medical condition but were not tagged as relevant in Gemma. We reviewed these datasets and judged whether there was a case for annotations to be added to Gemma. We concluded that there might be a case for nine of these datasets (Additional Data File 2). The Gemma team agreed that the “juvenile idiopathic arthritis” ontology term should be added for GSE13501. For GSE149632, we found that the “Parkinson disease” term had been added after we downloaded Gemma annotations and made our predictions. For 6 datasets, the Gemma curators agreed that the GEO series were relevant to the specified medical condition; however, these datasets were from cell lines. Current practice at Gemma is to annotate such datasets with the ontology term for the cell line; inferences regarding the associated medical conditions can be made via querying the respective ontologies. GSE8650 was tagged with “systemic juvenile idiopathic arthritis,” which is a distinct subtype of the juvenile idiopathic arthritis^54^. Researchers looking to study the condition more broadly might be interested in using this dataset, but they would need to perform ontology-based inference in Gemma to find it. For five additional datasets, we agreed that it was not appropriate to add annotations for the medical conditions we evaluated; however, the review process pointed to annotations that should be added or removed. For example, the language models suggested that GSE19697 was relevant to triple-negative breast carcinoma, yet the samples were from patients with basal-like breast carcinoma. These conditions have etiological overlap^55^, but they are distinct. The curators annotated the dataset in Gemma with the “basal-like breast carcinoma” term.

To gain additional insight into the top-performing model’s performance (`thenlper/gte-large`), we compared the length (number of characters after cleaning steps) of the dataset descriptions versus the cosine similarity scores for the GEO series in our test sets. We surmised that datasets with longer descriptions might be more likely to be ranked as relevant. However, we did *not* observe a significant correlation between these variables (Figure S2).

Finally, to evaluate the potential to find additional datasets relevant to these medical conditions, we used `thenlper/gte-large` to rank all non-Gemma GEO datasets that matched our filtering criteria. First, we created an average embedding for each medical condition, using descriptions from Gemma-annotated datasets as inputs. Second, we ranked each non-Gemma dataset based on the cosine similarity between its embedding and the averaged Gemma embedding for that condition. The Gemma curators reviewed the 50 datasets with the highest similarity per medical condition; these datasets were divided between the curators, with some overlap to support an evaluation of inter-rater agreement (Methods). The curators reviewed the dataset descriptions and selected the medical condition that they believed was most relevant. In cases where they lacked certainty, the reviewers indicated that “maybe” a dataset was relevant to a specified condition. If they believed none of the datasets was relevant, they selected “Other” (see Methods). We analyzed the datasets that the curators confidently classified as relevant to a particular medical condition. Taking into account both reviewers’ responses, the datasets selected by the language model were relevant 62-63% of the time, whereas only 50-53% of datasets selected by GEO’s search tool were relevant (Table 1; Additional Data File 3). These percentages differed considerably across the medical conditions (Figure 7). For Down syndrome, juvenile idiopathic arthritis, neuroblastoma, and triple-negative breast carcinoma, the language model produced more relevant results than GEO’s search tool. However, for Parkinson’s disease, GEO performed slightly better. For bipolar disorder, GEO returned only 21 results, and the curators agreed that these datasets were relevant 59.4% of the time. For the top-50 datasets returned by the language model, the curators agreed that the datasets were relevant only 21.7% of the time.

**Figure 7:**
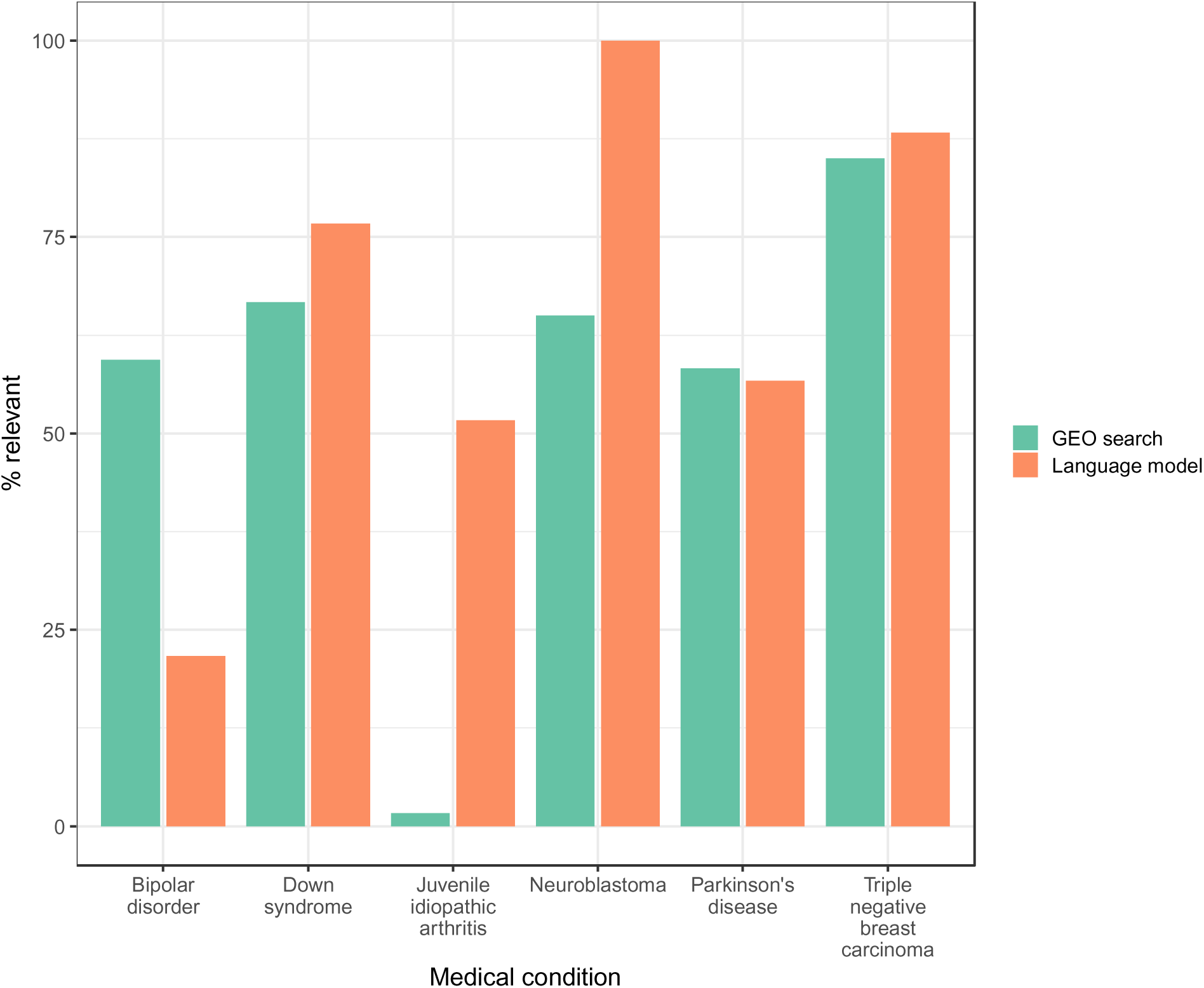
Agreement between curators’ responses and top search results for non-Gemma series. We reviewed the top series returned by GEO’s Advanced Search Builder or the `thenlper/gte-large` model for each medical condition. Curators manually reviewed the GEO descriptions and judged whether each series was relevant to a given medical condition or was “maybe” relevant. For bipolar disorder, the GEO search returned only 21 series.

**Table 1:** Comparisons of automated predictions with curators’ conclusions. This table indicates the number of times that each of two human curators indicated that a given GEO series was (or maybe was) relevant to one of six medical conditions for non-Gemma series. The percentages indicate the frequencies with which the medical condition predicted by the top-performing language model (`thenlper/gte-large`)—or GEO’s Advanced Search Builder—agreed with the medical condition selected by the human curators.

| Search method | Curator | # yes | % yes - relevant | # maybe | % maybe - relevant |
| --- | --- | --- | --- | --- | --- |
| Language model | 1 | 156 | 62.2 | 24 | 95.8 |
| Language model | 2 | 171 | 63.2 | 9 | 100.0 |
| GEO search | 1 | 139 | 50.4 | 27 | 92.6 |
| GEO search | 2 | 159 | 52.8 | 7 | 85.7 |

Some of the datasets categorized as irrelevant are instructive. For example, many predicted as relevant to bipolar disorder were instead found to pertain to schizophrenia or major depressive disorder; these conditions have overlapping features and shared genetic or neurobiological factors. Some predicted as relevant to juvenile idiopathic arthritis were focused on rheumatoid arthritis or pediatric systemic lupus erythematosus. These conditions share overlapping pathophysiological processes, but the curators did not categorize them as relevant because they are distinct conditions.

Of the 260 unique datasets returned by either search method that the curators deemed to be relevant, only 41 (15.8%) overlapped between the two methods. This finding suggests that the two search methods are complementary and thus can be used together.

Given the `thenlper/gte-large` model’s ability to identify relevant datasets, we created *GEOfinder*, a Web application that enables researchers to query GEO. We designed this application with the idea that researchers will first use GEO’s Advanced Search Builder to identify some datasets relevant to their research topic. When a researcher performs a search in GEO, it returns a list of datasets and provides an option to select a checkbox next to each dataset.

It also provides an option for users to download a file with information about the selected datasets. After completing these steps for datasets of interest, the researcher can upload the file into GEOfinder and search for datasets that have similar descriptions. GEOfinder uses language embeddings and cosine similarity to rank candidate datasets. GEOfinder can be found at https://bioapps.byu.edu/geofinder. Its source code is available from https://github.com/abbymuir47/GEOfinder3.0.

## Discussion

This project was motivated by our prior experiences searching for gene-expression datasets relevant to specific human diseases. Those efforts were time and labor intensive, and we sensed that we were missing many available datasets. GEO’s Advanced Search Builder supports queries based on keywords or keyphrases, accompanied by filtering options, whereas our study is based on the assumption that relatively long, narrative descriptions of datasets characterize their research context and thus should be more helpful for dataset finding. Even though individual researchers who deposit datasets vary widely in the ways they describe data and experiments, those researchers have domain expertise on the topic they are studying. Thus, we expect that the language they use generally reflects the biomedical context of each study and thus should be differentiable from language used in other contexts.

GEO does provide a collection of “DataSets” that have been human curated and annotated and thus are more easily searchable. However, as of January 13, 2025, only 4,348 Datasets had been curated, whereas 244,358 GEO series existed. Another option is MetaSRA, which contains annotations for 718,384 samples, most of which are from RNA-Sequencing experiments^19^. These annotations were generated using a semi-automated process; however, the last update occurred in 2020, and RNA-Sequencing experiments constitute only a portion of GEO. Although it would be preferable for human curators to annotate all GEO series and samples retroactively, this is unlikely, due to the time and resources that would be required. Therefore, we need methodologies that do not require manual annotation by experts. One innovation of our approach is that it essentially places a curation responsibility on the researchers who wish to find data. After they have identified *some* existing datasets that align with their research topic, the metadata from those datasets serve as “ground truth” examples in the search for additional ones.

Our goal was not necessarily to supplant existing search tools, which are effective in many cases and are often ontology backed. Nor do we claim that one or a few language models are better than all others. It would be infeasible to make this claim due to the massive number of available models and the rapid pace at which modeling methodologies and training corpora are changing. Rather, our goals were 1) to explore the possibility of using language models to aid with dataset finding, 2) to evaluate how much the models would differ from each other in their performance, 3) to assess how well they would perform in different biomedical contexts, and 4) to compare them against GEO’s Advanced Search Builder. In doing so, we found that the language models often returned very different results than GEO’s tool, thus supporting the idea that these approaches are complementary.

Some models performed quite well and dramatically better than others. Generally, models with larger embedding sizes performed better than those with smaller sizes, but this observation is confounded because embedding sizes have co-evolved with modeling methodologies and data sources. It is possible that with additional optimizations—such as using a different chunking strategy or fine tuning based on human feedback—the model rankings would change considerably. Performance also varied according to the medical condition being studied. One might presume that increasing the number of examples in a reference set would lead to better precision and recall. However, our methodology achieved excellent performance for juvenile idiopathic arthritis, the condition with the fewest examples of the six we studied. This suggests that accurately identifying new datasets may depend less on the number of examples and more on the distinctiveness of the language used to describe a particular medical condition.

Our study was limited to human data and datasets related to medical research. Currently, the extent to which our findings generalize to other contexts is unclear. An additional limitation is that the manual reviews were somewhat subjective. We conceptualized medical relevance as whether researchers studying a given medical condition might be interested in using the data to study that condition (Additional Data File 1). However, it was sometimes difficult to make this distinction. For example, when evaluating neuroblastoma, a form of cancer that affects nerve cells, we identified many GEO series that had used neuroblastoma cell lines but not necessarily to study neuroblastoma itself. For example, Schartner, et al used the SH-SY5Y cell line to study the downstream effects of a single-nucleotide variant, irrespective of any medical condition^56^. Fardin, et al used neuroblastoma cell lines to study differences between normoxic and hypoxic conditions; they also evaluated benefits of using a particular algorithm for detecting relevant genes^57^. Although their overarching goal was to shed light on neuroblastoma development, that connection was indirect for this study.

Although our findings are promising, there are opportunities for further exploration. Our method uses only high-level dataset descriptions as inputs. In contrast, PEPHub^27^ uses sample-level metadata to construct embeddings. The molecular data from these studies may also provide useful information. Combining some or all of these information types may yield better results for dataset finding. When searching for semantically similar datasets, we identified multiple datasets known to be relevant and averaged their embeddings before searching for other datasets with similar embeddings. A better alternative might be to keep the embeddings separate and train a classification algorithm to differentiate between relevant and irrelevant examples.

In conclusion, our findings provide strong evidence that language models hold promise for enhancing retrieval of publicly available datasets. Our evaluation, which included a diverse set of models, offers an impartial comparison, as we did not contribute to the development of any models under investigation. Yet despite the effectiveness of many models in our evaluation, it is clear that retrieval performance needs further improvement and that human reviews remain essential for the foreseeable future. While our study specifically targeted gene-expression data, the approach we developed can be applied to other types of molecular data that are accompanied by human-language descriptions. We hope that these efforts lead to improvements in data accessibility, thus increasing the value derived from previous research investments.

## Additional Data Files

**Additional Data File 1: Rubric that the curators used to evaluate whether GEO datasets were relevant to the medical conditions.**

**Additional Data File 2: GEO series most frequently predicted to be associated with a particular medical condition but that were not annotated as such in GEO.** We reviewed each series and recorded a comment justifying our conclusion about whether there was a strong case to add an annotation in Gemma for each series.

**Additional Data File 3: Top-ranked datasets with curator responses** Using the `thenlper/gte-large` model, we ranked the non-Gemma GEO datasets based on semantic similarity to the Gemma datasets annotated for the six medical conditions we studied. Additionally, we used GEO’s Advanced Search builder to search for datasets associated with each medical condition. Two curators reviewed these results and indicated their findings in this spreadsheet.

## References

1. Wilkinson, M. D. et al. The FAIR Guiding Principles for scientific data management and stewardship. Scientific Data 3, 160018 (2016).

2. Clough, E. & Barrett, T. The Gene Expression Omnibus Database. Methods Mol Biol 1418, 93–110 (2016).

3. Lipscomb, C. E. Medical subject headings (MeSH). Bulletin of the Medical Library Association 88, 265 (2000).

4. Gendoo, D. M. et al. MetaGxData: Clinically annotated breast, ovarian and pancreatic cancer datasets and their use in generating a multi-cancer gene signature. Scientific Reports 9, 8770 (2019).

5. Ganzfried, B. F. et al. curatedOvarianData: Clinically annotated data for the ovarian cancer transcriptome. Database (Oxford) 2013, bat013 (2013).

6. Zimmermann, P. et al. ExpressionData - A public resource of high quality curated datasets representing gene expression across anatomy, development and experimental conditions. BioData Mining 7, 18 (2014).

7. Villaseñor-Altamirano, A. B. et al. PulmonDB: A curated lung disease gene expression database. Sci Rep 10, 514 (2020).

8. Federico, A. et al. Manually curated and harmonised transcriptomics datasets of psoriasis and atopic dermatitis patients. Sci Data 7, 343 (2020).

9. Rahman, M. et al. A curated transcriptome dataset collection to investigate the functional programming of human hematopoietic cells in early life. F1000Res 5, 414 (2016).

10. Lim, N. et al. Curation of over 10 000 transcriptomic studies to enable data reuse. Database 2021, baab006 (2021).

11. Wang, Z. et al. Extraction and analysis of signatures from the Gene Expression Omnibus by the crowd. Nature Communications 7, 12846 (2016).

12. Hadley, D. et al. Precision annotation of digital samples in NCBI’s gene expression omnibus. Sci Data 4, 170125 (2017).

13. Lachmann, A. et al. Massive mining of publicly available RNA-seq data from human and mouse. Nature Communications 9, 1366 (2018).

14. Shah, N. et al. A crowdsourcing approach for reusing and meta-analyzing gene expression data. Nature Biotechnology 34, 803–806 (2016).

15. Good, B. M. & Su, A. I. Crowdsourcing for bioinformatics. Bioinformatics 29, 1925– 1933 (2013).

16. Perez-Riverol, Y. et al. Quantifying the impact of public omics data. Nat Commun 10, 3512 (2019).

17. Ohno-Machado, L. et al. Finding useful data across multiple biomedical data repositories using DataMed. Nature Genetics 49, 816–819 (2017).

18. Chen, X. et al. DataMed – an open source discovery index for finding biomedical datasets. Journal of the American Medical Informatics Association 25, 300–308 (2018).

19. Bernstein, M. N., Doan, A. & Dewey, C. N. MetaSRA: Normalized human sample-specific metadata for the Sequence Read Archive. Bioinformatics 33, 2914–2923 (2017).

20. Hawkins, N. T., Maldaver, M., Yannakopoulos, A., Guare, L. A. & Krishnan, A. Systematic tissue annotations of genomics samples by modeling unstructured metadata. Nat Commun 13, 6736 (2022).

21. Alameer, A. & Chicco, D. geoCancerPrognosticDatasetsRetriever: A bioinformatics tool to easily identify cancer prognostic datasets on Gene Expression Omnibus (GEO). Bioinformatics 38, 1761–1763 (2022).

22. Chen, G. et al. Restructured GEO: Restructuring Gene Expression Omnibus metadata for genome dynamics analysis. Database 2019, bay145 (2019).

23. Djordjevic, D. et al. Discovery of perturbation gene targets via free text metadata mining in Gene Expression Omnibus. Computational Biology and Chemistry 80, 152–158 (2019).

24. Chua, H.-E., Tucker-Kellogg, L. & Bhowmick, S. S. ArcheGEO: Towards improving relevance of gene expression omnibus search results. in Proceedings of the 13th ACM International Conference on Bioinformatics, Computational Biology and Health Informatics 1– 10 (Association for Computing Machinery, New York, NY, USA, 2022). doi:10.1145/3535508.3545531.

25. Giles, C. B. et al. ALE: Automated label extraction from GEO metadata. BMC bioinformatics 18, 7–16 (2017).

26. Lake, B. M. & Murphy, G. L. Word meaning in minds and machines. Psychological Review 130, 401–431 (2023).

27. LeRoy, N. J. et al. PEPhub: A database, web interface, and API for editing, sharing, and validating biological sample metadata. GigaScience 13, giae033 (2024).

28. Patra, B. G., Roberts, K. & Wu, H. A content-based dataset recommendation system for researchers—a case study on Gene Expression Omnibus (GEO) repository. Database 2020, baaa064 (2020).

29. Mikolov, T., Chen, K., Corrado, G. & Dean, J. Efficient estimation of word representations in vector space. *arXiv preprint arXiv:1301*.3781 (2013).

30. Pennington, J., Socher, R. & Manning, C. D. Glove: Global vectors for word representation. in Proceedings of the 2014 conference on empirical methods in natural language processing (EMNLP) 1532–1543 (2014).

31. Devlin, J. Bert: Pre-training of deep bidirectional transformers for language understanding. arXiv preprint arXiv:*1810.04805* (2018).

32. Wolf, T. et al. Transformers: State-of-the-Art Natural Language Processing. In Proceedings of the 2020 Conference on Empirical Methods in Natural Language Processing: System Demonstrations (eds. Liu, Q. & Schlangen, D.) 38–45 (Association for Computational Linguistics, Online, 2020). doi:10.18653/v1/2020.emnlp-demos.6.

33. Harnoune, A. et al. BERT based clinical knowledge extraction for biomedical knowledge graph construction and analysis. Computer Methods and Programs in Biomedicine Update 1, 100042 (2021).

34. Lee, J. et al. BioBERT: A pre-trained biomedical language representation model for biomedical text mining. Bioinformatics 36, 1234–1240 (2020).

35. Chen, Q. et al. BioConceptVec: Creating and evaluating literature-based biomedical concept embeddings on a large scale. PLoS Comput Biol 16, e1007617 (2020).

36. Remy, F., Demuynck, K. & Demeester, T. BioLORD-2023: Semantic textual representations fusing large language models and clinical knowledge graph insights. Journal of the American Medical Informatics Association 31, 1844–1855 (2024).

37. Alsentzer, E., et al. Publicly Available Clinical BERT Embeddings. (2019) doi:10.48550/arXiv.1904.03323.

38. Chiu, B., Crichton, G., Korhonen, A. & Pyysalo, S. How to Train good Word Embeddings for Biomedical NLP. in *Proceedings of the 15th Workshop on Biomedical Natural Language Processing* 166–174 (Association for Computational Linguistics, Berlin, Germany, 2016). doi:10.18653/v1/W16-2922.

39. Wang, Y. et al. A comparison of word embeddings for the biomedical natural language processing. Journal of Biomedical Informatics 87, 12–20 (2018).

40. Khoroshevskyi, O., LeRoy, N., Reuter, V. P. & Sheffield, N. C. GEOfetch: A command-line tool for downloading data and standardized metadata from GEO and SRA. Bioinformatics 39, btad069 (2023).

41. Richardson, L. Beautiful soup documentation. (2007).

42. Bird, S., Klein, E. & Loper, E. Natural Language Processing with Python: Analyzing Text with the Natural Language Toolkit. (" O’Reilly Media, Inc.", 2009).

43. Vasilevsky, N. et al. Mondo Disease Ontology: Harmonizing disease concepts across the world. in CEUR-WS vol. 2807 (2020).

44. Noy, N. F. et al. BioPortal: Ontologies and integrated data resources at the click of a mouse. Nucleic Acids Res 37, W170–W173 (2009).

45. Lowe, H. J. & Barnett, G. O. Understanding and using the medical subject headings (MeSH) vocabulary to perform literature searches. Jama 271, 1103–1108 (1994).

46. Wolf, T. et al. HuggingFace’s Transformers: State-of-the-art Natural Language Processing. (2020) doi:10.48550/arXiv.1910.03771.

47. Bojanowski, P., Grave, E., Joulin, A. & Mikolov, T. Enriching word vectors with subword information. *arXiv preprint arXiv:1607.04606* (2016).

48. Mikolov, T., Grave, E., Bojanowski, P., Puhrsch, C. & Joulin, A. Advances in pre-training distributed word representations. in Proceedings of the international conference on language resources and evaluation (LREC 2018) (2018).

49. OpenAI. Openai: The official Python library for the openai API.

50. Wickham, H. et al. Welcome to the tidyverse. Journal of Open Source Software 4, 1686 (2019).

51. Piccolo, S. R. & Frampton, M. B. Tools and techniques for computational reproducibility. Gigascience 5, 30 (2016).

52. Cao, H. Recent advances in text embedding: A Comprehensive Review of Top-Performing Methods on the MTEB Benchmark. (2024) doi:10.48550/arXiv.2406.01607.

53. Liu, Y., et al. Roberta: A robustly optimized bert pretraining approach. arXiv preprint arXiv:1907.11692 (2019).

54. Mellins, E. D., Macaubas, C. & Grom, A. A. Pathogenesis of systemic juvenile idiopathic arthritis: Some answers, more questions. Nat Rev Rheumatol 7, 416–426 (2011).

55. Seal, M. D. & Chia, S. K. What Is the Difference Between Triple-Negative and Basal Breast Cancers? The Cancer Journal 16, 12 (2010-01/2010-02).

56. Schartner, C. et al. The regulation of tetraspanin 8 gene expression—A potential new mechanism in the pathogenesis of bipolar disorder. American Journal of Medical Genetics Part B: Neuropsychiatric Genetics 174, 740–750 (2017).

57. Fardin, P. et al. The L1-L2 regularization framework unmasks the hypoxia signature hidden in the transcriptome of a set of heterogeneous neuroblastoma cell lines. BMC Genomics 10, 474 (2009).

58. Brand, A., Allen, L., Altman, M., Hlava, M. & Scott, J. Beyond authorship: Attribution, contribution, collaboration, and credit. Learned Publishing 28, 151–155 (2015).

