## Supplementary Material for "Using semantic search to find publicly available gene-expression datasets"

### Supplementary Tables

**Table S1: Language model characteristics.** This table lists the models that we compared. The first part of each model name indicates the organization, person, or package who created the model. The second part of each model name is a unique identifier that describes the methodology and/or data source(s) used for the model. The embedding size indicates the number of dimensions in the vector space. The remaining columns indicate which type of corpus was used for training, the general type of language model that was trained, and how fine tuning was performed (when applicable).

| **Model** | **Embedding size** | **Data source type** | **Model category** | **Fine tuning** |
| --- | --- | --- | --- | --- |
| FremyCompany/BioLORD-2023 | 768 | Biomedical | Ontology-driven concept representations | This model is based on sentence-transformers/all-mpnet-base-v2 and was further fine tuned on the BioLORD-Dataset and LLM-generated definitions from the Automatic Glossary of Clinical Terminology (AGCT). |
| NeuML/pubmedbert-base-embeddings | 768 | Biomedical | Bidirectional Encoder Representations from Transformers (BERT) |  |
| albert/albert-base-v2 | 768 | General | Masked language modeling (MLM) |  |
| albert/albert-xxlarge-v2 | 4096 | General | Masked language modeling (MLM) | This checkpoint ostensibly is a fine-tuning of albert/albert-xxlarge-v2. |
| allenai/biomed_roberta_base | 768 | Biomedical | Bidirectional Encoder Representations from Transformers (BERT) |  |
| allenai/scibert_scivocab_uncased | 768 | Scientific | Bidirectional Encoder Representations from Transformers (BERT) |  |
| emilyalsentzer/Bio_ClinicalBERT | 768 | Biomedical | Bidirectional Encoder Representations from Transformers (BERT) |  |
| fasttext/cbow-commoncrawl | 300 | General | Continuous Bag of Words (Word2Vec) |  |
| fasttext/cbow-wikinews | 300 | General | Continuous Bag of Words (Word2Vec) |  |
| hkunlp/instructor-xl | 768 | General | Instruction-fine-tuned text embedding model |  |
| medicalai/ClinicalBERT | 768 | Biomedical | Bidirectional Encoder Representations from Transformers (BERT) |  |
| microsoft/BiomedNLP-BiomedBERT-base-uncased-abstract-fulltext | 768 | Biomedical | Bidirectional Encoder Representations from Transformers (BERT) |  |
| nomic-ai/nomic-embed-text-v1.5 | 768 | General | Matryoshka Representation Learning |  |
| nuvocare/WikiMedical_sent_biobert | 768 | Biomedical | Bidirectional Encoder Representations from Transformers (BERT) | Based on the dmis-lab/biobert-base-cased-v1.2 backbone and has been trained on the WikiMedical_sentence_simialrity dataset. |
| openai/text-embedding-3-large | 3072 | General | Generative Pre-trained Transformer (GPT) |  |
| openai/text-embedding-3-small | 1536 | General | Generative Pre-trained Transformer (GPT) |  |
| openai/text-embedding-ada-002 | 1536 | General | Generative Pre-trained Transformer (GPT) |  |
| pritamdeka/S-BioBert-snli-multinli-stsb | 768 | Biomedical | Bidirectional Encoder Representations from Transformers (BERT) |  |
| pritamdeka/S-Biomed-Roberta-snli-multinli-stsb | 768 | Biomedical | Bidirectional Encoder Representations from Transformers (BERT) | The base model used is allenai/biomed_roberta_base which has been fine-tuned for sentence similarity. |
| pritamdeka/S-PubMedBert-MS-MARCO-SCIFACT | 768 | Biomedical | Bidirectional Encoder Representations from Transformers (BERT) |  |
| sentence-transformers/all-MiniLM-L6-v2 | 384 | General | Self-supervised contrastive learning | We used the pretrained nreimers/MiniLM-L6-H384-uncased model and fine-tuned in on a 1B sentence pairs dataset. |
| sentence-transformers/all-mpnet-base-v2 | 768 | General | Self-supervised contrastive learning | We used the pretrained microsoft/mpnet-base model and fine-tuned in on a 1B sentence pairs dataset. |
| sentence-transformers/all-roberta-large-v1 | 1024 | General | Self-supervised contrastive learning | We used the pretrained roberta-large model and fine-tuned in on a 1B sentence pairs dataset. |
| sentence-transformers/average_word_embeddings_glove.6B.300d | 300 | General | Global Vectors for Word Representation (GloVe) |  |
| sentence-transformers/average_word_embeddings_glove.840B.300d | 300 | General | Global Vectors for Word Representation (GloVe) |  |
| sentence-transformers/msmarco-distilbert-base-v3 | 768 | General | Bidirectional Encoder Representations from Transformers (BERT) |  |
| sentence-transformers/paraphrase-TinyBERT-L6-v2 | 768 | General | Bidirectional Encoder Representations from Transformers (BERT) |  |
| sentence-transformers/sentence-t5-large | 768 | General | Text-to-text transformers (T5) |  |
| sentence-transformers/sentence-t5-xl | 768 | General | Text-to-text transformers (T5) |  |
| thenlper/gte-large | 1024 | General | Multi-stage contrastive learning |  |

**Table S2: Performance when chunking longer texts versus not chunking longer texts.** When input texts are relatively long, they can be broken into smaller chunks. An embedding is derived for each chunk, and an embedding is calculated for each input text as the average of the individual-chunk embeddings. This table shows the median increase or decrease (across six medical conditions) in the area under the precision-recall curve (AUPRC) when chunking *was* used, relative to the AUPRC when chunking was *not* used.

| **Method** | **Difference** |
| --- | --- |
| albert/albert-base-v2 | -0.0021747 |
| albert/albert-xxlarge-v2 | 0.0016946 |
| allenai/biomed_roberta_base | 0.0020904 |
| allenai/scibert_scivocab_uncased | -0.0029334 |
| emilyalsentzer/Bio_ClinicalBERT | 0.0017210 |
| fasttext/cbow-commoncrawl | 0.0000000 |
| fasttext/cbow-wikinews | 0.0000000 |
| FremyCompany/BioLORD-2023 | -0.2920301 |
| hkunlp/instructor-xl | -0.1756387 |
| medicalai/ClinicalBERT | -0.0158756 |
| microsoft/BiomedNLP-BiomedBERT-base-uncased-abstract-fulltext | -0.0207285 |
| NeuML/pubmedbert-base-embeddings | -0.2329609 |
| nomic-ai/nomic-embed-text-v1.5 | -0.4027749 |
| nuvocare/WikiMedical_sent_biobert | -0.3032933 |
| openai/text-embedding-3-large | -0.2633428 |
| openai/text-embedding-3-small | -0.3064818 |
| openai/text-embedding-ada-002 | -0.3695718 |
| pritamdeka/S-BioBert-snli-multinli-stsb | -0.2239655 |
| pritamdeka/S-Biomed-Roberta-snli-multinli-stsb | -0.1745970 |
| pritamdeka/S-PubMedBert-MS-MARCO-SCIFACT | -0.0404077 |
| sentence-transformers/all-MiniLM-L6-v2 | -0.2525039 |
| sentence-transformers/all-mpnet-base-v2 | -0.2652256 |
| sentence-transformers/all-roberta-large-v1 | -0.4321781 |
| sentence-transformers/average_word_embeddings_glove.6B.300d | -0.0062240 |
| sentence-transformers/average_word_embeddings_glove.840B.300d | -0.0029985 |
| sentence-transformers/msmarco-distilbert-base-v3 | -0.1799446 |
| sentence-transformers/paraphrase-TinyBERT-L6-v2 | -0.3346350 |
| sentence-transformers/sentence-t5-large | -0.2030918 |
| sentence-transformers/sentence-t5-xl | -0.1469890 |
| thenlper/gte-large | -0.4557162 |

### Supplementary Figures

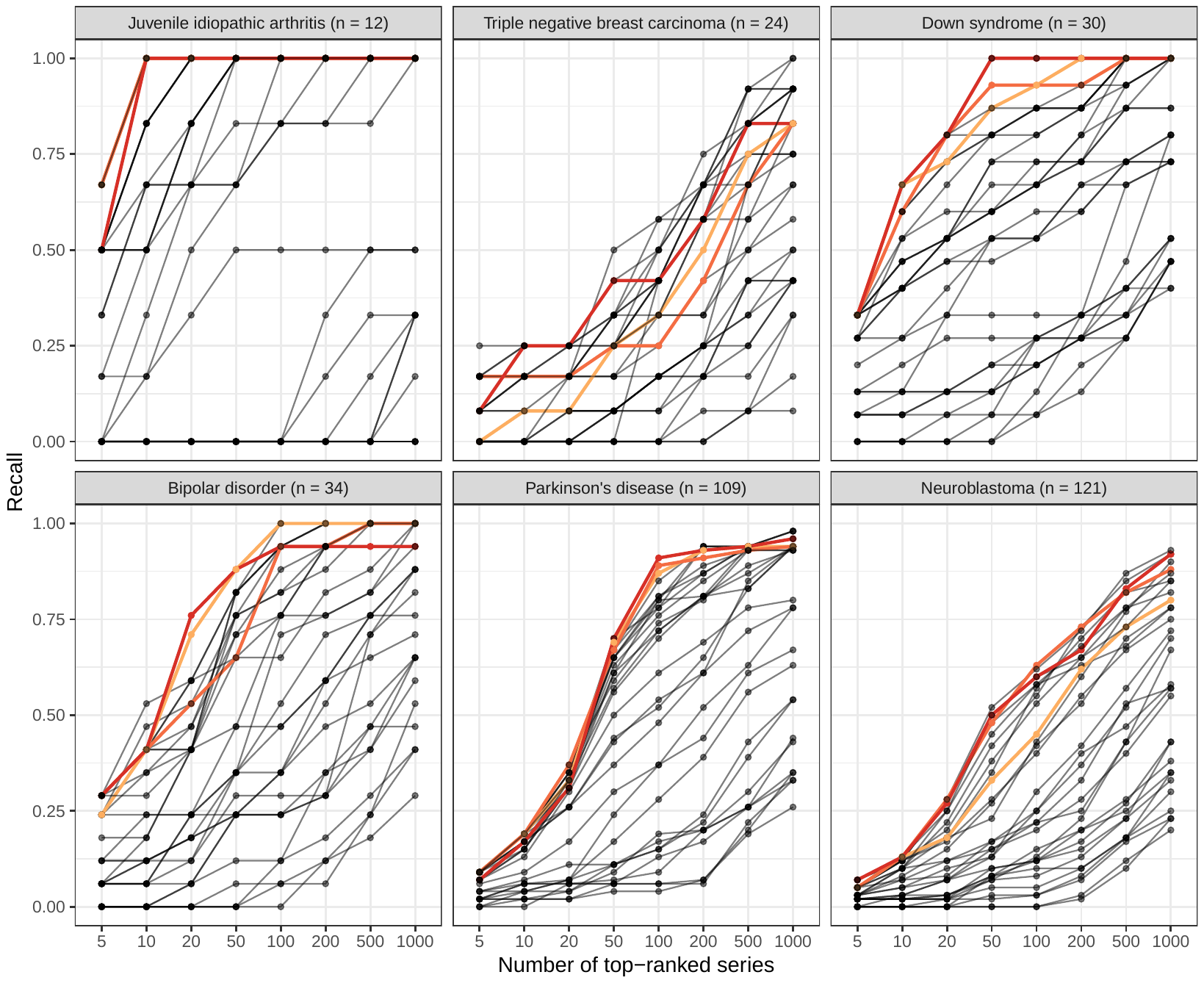

**Figure S1: Within-Gemma recall (sensitivity) for top-*n* ranked results returned from language models.** We used language models to identify Gemma datasets that had been annotated as relevant to six medical conditions. For each medical condition and language model, we calculated the proportion of relevant datasets that ranked among the top-*n* results. A recall value of 1.0 indicates that all Gemma-annotated datasets were included in the top-*n* results.

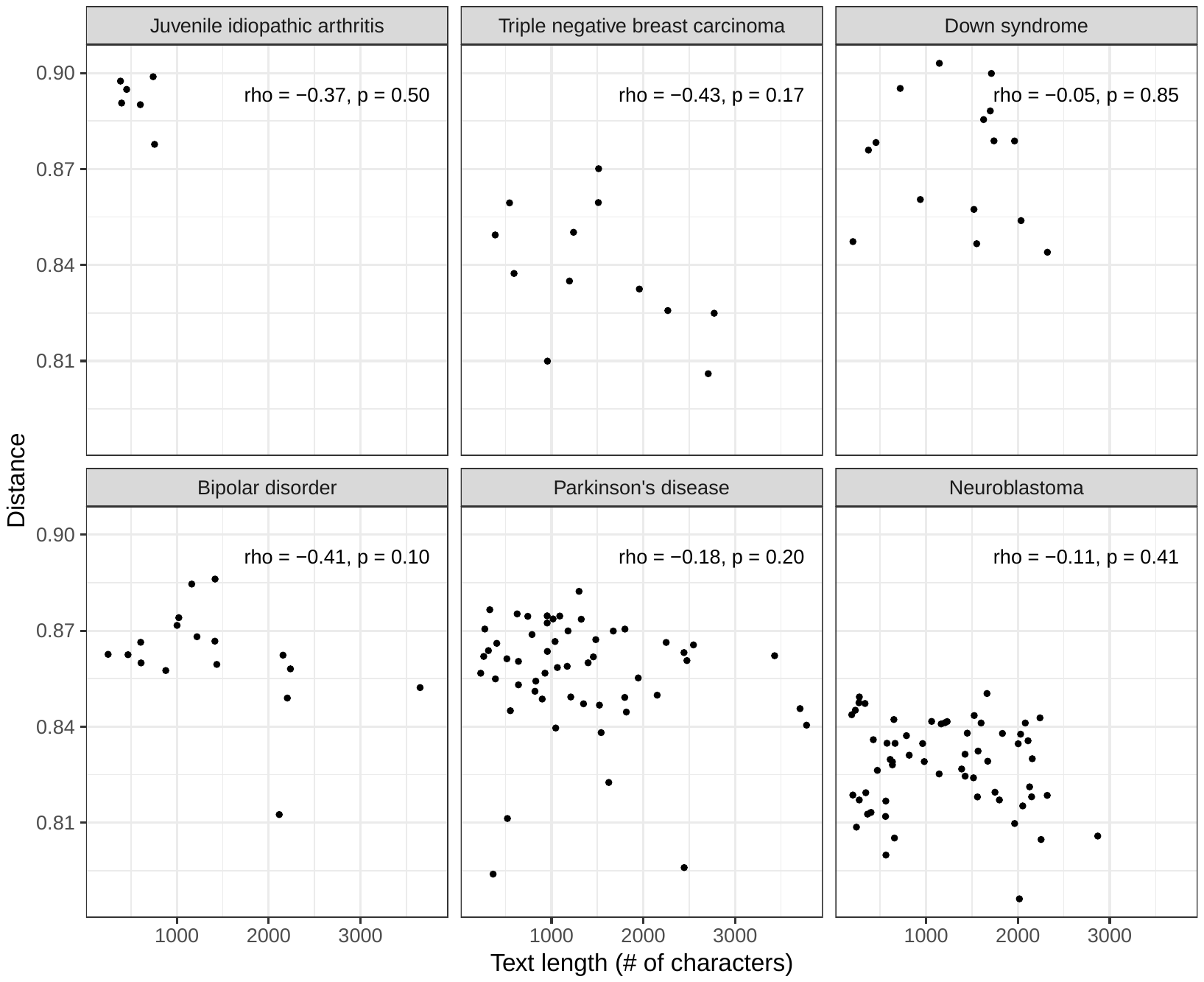

**Figure S2: Comparison between the length of series descriptions and relevance rankings.** We calculated the length (number of characters after cleaning steps) of the GEO series descriptions and the order in which the thenlper/gte-large model ranked the series. Spearman’s rank correlation test was used to compare these variables for each medical condition.
