## Additional Data File 1 for "Using semantic search to find publicly available gene-expression datasets"

In the spreadsheet, each row represents a GEO series. You will find a title, description, and a summary of the overall study design for each series. Additionally, you will see columns for each of six medical conditions. Review the information for each GEO series. Put "Yes," "No," or "Maybe" in the cells for the medical conditions. But only put "Yes" (or "Maybe") in one of those cells. The remaining cells should have "No". (We have filled these with "No" as defaults to reduce the amount of typing you have to do.) Only use the information provided in the spreadsheet. However, if you find that more investigation (such as looking at a journal article associated with the GEO series) would be helpful, indicate that in the "Investigate more" column of the spreadsheet; we anticipate that this will not be needed in most cases. You can also add comments under "Optional comments."

We conceptualize medical relevance as whether a study uses samples from people with the medical condition AND whether the researchers were studying that medical condition. Some studies use cell lines as samples (such as MDA-MB-257 or IMR-32). Resources like Cellosaurous and the Cell Line Ontology may help with identifying the medical condition associated with particular cell lines. Sometimes the GEO descriptions use generic terms. For example, a description might say "breast carcinoma." You should only indicate that study as relevant if it specifically had samples from triple-negative breast carcinomas (this is just one example), not from a more general disease. Also keep in mind that terms differ. For example, "carcinoma" might be used by one researcher, whereas "cancer" or "tumor" might be used by another. Use the synonyms in the Mondo Disease Ontology to clarify, as needed.

It is fine if a dataset contains samples from multiple medical conditions. As long as there are at least some samples associated with one of the medical conditions, mark it as "Yes."
